# Spatiotemporal Gray Matter Plasticity During Chronification of Preclinical Neuropathic Pain

**DOI:** 10.1101/2025.10.03.679890

**Authors:** Alexis Levesque, Ines Charef, Marc-Antoine Fortier, Sabrina Dumont, Natalia Muñoz Gómez, Marc-André Dansereau, Gabriel Richard, Pascal Tétreault

## Abstract

Chronic neuropathic pain is increasingly recognized as a brain disease characterized by time-dependent structural and functional reorganization of key neural circuits. While human imaging studies implicate widespread changes in network connectivity and gray matter density (GMD), animal models enable direct longitudinal mapping of such plasticity. Here, we applied high-resolution structural MRI in a rat model of chronic pain (spared nerve injury, SNI) and quantified GMD changes across 134 brain regions. Dynamic weight bearing analysis confirmed persistent pain in SNI rats, validating the chronic pain phenotype in our experimental cohort. Longitudinal MRI revealed significant GMD alterations in 31 regions, predominantly within limbic, prefrontal, and cingulate circuits, representing 21% of total brain volume. Among this affected volume, over 17% of brain volume demonstrated GMD increases while only ~3% showed GMD decreases, indicating a heterogeneous neuroplastic response. Specifically, the Frontal Association Cortex exhibited an approximate 10% increase in GMD, the Primary Cingular Cortex showed a modest increase of about 2%, and the Amygdalohyppocampic Area demonstrated a ~10% decrease in GMD over 28 days. Primary sensory, parietal, visual, retrosplenial, and temporal cortices remained largely unaffected. No significant changes were observed in healthy animals over the same period, highlighting the specificity of brain reorganization to persistent neuropathic pain. These findings reaffirm the ability of MRI to robustly quantify pain-induced neuroanatomical remodeling but leave open critical questions about the underlying cellular and molecular mechanisms. Future studies integrating histological and molecular approaches are needed to determine the precise substrate and reversibility of these structural changes, with the goal of identifying therapeutic targets to prevent or reverse maladaptive neuroplasticity in chronic.

## Introduction

Chronic neuropathic pain is increasingly recognized as a brain disease, characterized not only by continual nociceptive input but also by profound, time-dependent reorganization of brain networks at both functional and structural levels^1,2^. Human neuroimaging research robustly links chronic pain to widespread changes in network connectivity and gray matter architecture, particularly within sensory, limbic, and cognitive control circuits^3–5^. Such changes often manifest as reductions in gray matter density (GMD) and volume in prefrontal, insular, cingulate, hippocampal, and striatal regions, areas repeatedly implicated in pain processing, emotion, and executive function^6–8^. Critically, these neuroplastic changes are thought not just to reflect consequences of chronic pain but to participate in its maintenance and chronification^1,9–11^.

However, there is a persistent and important gap in our understanding of the nature, timing, and significance of these structural changes, in large part because most human studies are cross-sectional and thus cannot disambiguate cause from consequence^1,11^. Though some clinical data suggest reversibility of cortical atrophy after pain relief^4^, preclinical longitudinal imaging is a powerful tool to clarify the trajectory and mechanistic underpinnings of these phenomena^12^.

Among animal models, the spared nerve injury (SNI) model in rats is a gold standard for preclinical research, enabling high-resolution, time-resolved investigation of both behavior and brain^9,13,14^. Landmark animal study by Seminowicz et al.^9^ demonstrated late decreases in cortical volume measured by MRI, specifically in the prefrontal cortex, retrosplenial, and entorhinal cortex volumes after SNI, temporally coinciding with the development of anxiety-like behaviors. Yet, most follow-up studies, while leveraging modalities such as fMRI, spectroscopy, and PET have focused on functional or metabolic changes, leaving the anatomical dimension underexplored^10,13–17^. For example, in rats, Hubbard et al.^13^ detailed early and late changes in brain metabolism and network activity associated with neuropathic pain behaviors but offered little insight into regional structural remodeling. Thompson et al.^14^ used PET imaging in animals to reveal persistent S1 hyperactivity, presumed to underlie ongoing pain, yet did not examine if functional changes accompany morphometric abnormalities.

In parallel, human functional MRI studies further clarify that, as neuropathic pain becomes chronic, brain activity shifts from classical sensory pathways toward circuits regulating affect and motivation, especially the limbic system (hippocampus, amygdala, nucleus accumbens (NAc), prefrontal and cingulate cortices)^10,15,18–20^. This pattern mirrors findings in chronic low back pain (CLBP) in human patients, where a reorganization of brain activity away from somatosensory toward limbic and prefrontal regions has also been demonstrated^21^. Important work by Baliki et al.^10^ and Chang et al.^15,19^ provides strong evidence that these limbic/motivation circuits orchestrate the transition to pain chronicity, with reorganization most evident after several weeks of ongoing pain. Wei et al.^20^ compellingly demonstrate that activating the dorsal hippocampus not only relieves neuropathic pain but also normalizes specific network connectivities again, with the structural basis of these changes largely unknown.

Moreover, molecular and pharmacological manipulation of the pain network (e.g. anti-NGF therapy)^22^ or sex and age modulations^18,23,24^ have pinpointed key modulators of functional network states and pain vulnerability in animal models. Yet, none of these modern studies systematically assessed concurrent changes in grey matter density, regional volume, or morphometry in the SNI model. Given that several clinical and multi-modal meta-analyses highlight the diagnostic power of brain morphometric signatures in chronic pain states^1,11^, this gap should be validated at the preclinical level. Thus, the field stands at an important point where the lack of longitudinal, regionally resolved mapping of brain morphometry in the SNI model^13,25,26^ needs to be addressed. Our study addresses these gaps by applying high-resolution, longitudinal structural MRI in the rat SNI model to systematically map regional changes in GMD over time, with a special focus on limbic, prefrontal, cingulate, and reward circuits highlighted by recent functional and pharmacological studies^9,10,13,15,18–20^.

## Methods

### Animals and surgical procedures

Adult male Sprague-Dawley rats (175-200g, 2-mo-old, Charles River, QC) were housed in pairs in standard shoebox cages connected to a ventilation rack, in a temperature-controlled (23 ± 1°C) environment (12:12 hrs light/dark cycle). The rodents had *ad lib* access to tap water and food. Animals were randomly assigned to either the SNI (*n*=9) or control (*n*=6) group. SNI involves the transection of two of the three distal branches of the sciatic nerve (tibial and peroneal) while “sparing” the sural nerve^27^. All experiments were carried out under the approval of the institutional ethic review board (2021-3227).

### Dynamic Weight Bearing assessment

Dynamic weight bearing (DWB) was assessed at baseline (the day before surgery) and on days 7, 14, 21, and 28 post-surgery. Prior to each testing session, animals were acclimated to the testing rooms for 20–30 minutes before being evaluated with the DWB device (Bioseb, Boulogne, France), which quantifies non-evoked pain and weight distribution in freely moving rats, as previously described^28^. The apparatus consists of a Plexiglass enclosure (22 × 22 × 30 cm) equipped with a floor sensor composed of 44 × 44 captors (10.89mm^2^ per captor) and a side-mounted camera. During a 5-minute session, rats were placed individually in the enclosure without prior acclimatization to maximize exploratory behaviors. Pressure data and video recordings were transmitted in real time to a laptop *via* a universal serial bus (USB) interface and processed using the DWB software v1.3. A zone was considered valid when at least one captor recorded ≥ 4 g and a minimum of two adjacent captors recorded ≥ 1 g. For reach time segment where weight distribution remained stable for more than 0.5 seconds, validated zones were assigned to specific paws or other body parts (e.g., tails, testicles) by and observer, based on the video and the activation map. For each animal, the mean percentage of body weight borne by each limb, particularly the ipsilateral (operated) hind paw, was calculated over the entire session. This measure allowed us to objectively confirm the presence of pain-related behavior, as animals with neuropathic pain typically reduce weight bearing on the affected limb.

### MRI acquisition & image analysis

Anesthesia was induced with 3-4% of isoflurane in 1.5 L/min of O2. Animals were scanned for 30 min using a Varian 7T small animal MRI scanner and a rat head surface coil (Rapid biomedical). The imaging protocol consisted of a scout image for localization and a single 3D gradient echo sequence covering the entire brain with the following parameters: TR 50 ms, TE 25 ms, flip angle 15°, 2 averages, matrix size of 192 × 192 × 96 voxel, field of view of 30 × 30 ×15 mm. The respiratory rate was monitored throughout the scan, and isoflurane was adjusted (range 1.5-3% in 1.5 L/min O2) to maintain a rate of approximately 60 breaths/min. Animals were kept warm using hot air to maintain an ambient temperature of 30 °C.

### MRI Processing

Images were processed using Python (v. 3.8), 3D Slicer (v. 5.0.3) and SPM12. First, images were converted from FDF to NIfTI using an in-house Python script. Images were then realigned with the 90 × 90 × 90 µm^3^ ex vivo SIGMA atlas^29^ in 3D Slicer using the BRAINS module. A scaling factor of 10 was applied to the voxel size of the atlas, tissue templates and individual images before segmentation in SPM12 to match the human brain size expected by the software. This method was compared with different adjustments to the default parameters to account for a smaller voxel size and was found to provide more accurate tissue segmentation.

To enable comparison with other studies and facilitate cross-atlas analyses, all images were also registered to the 39 × 39 × 39 µm^3^ ex vivo Waxholm Space (WHS) rat brain atlas^30^ v4.01^31^. Linear registration from the SIGMA space to the WHS was performed using ANTsPy (v. 0.5.3). The WHS atlas, in its original form, does not provide separate labels for left and right hemispheres. Therefore, we generated hemisphere-specific versions by following a multi-step process. First, we created masks for the left and reight hemispheres using the SIGMA atlas. Second, these masks were registered to the Waxholm atlas space with linear registration in ANTs. Third, the resulting masks were manually edited in ITK-snap to ensure that all brain regions were adequately covered. Finally, the masks were applied to the Waxholm atlas to generate right and left hemisphere regions, and region numbers were edited so that all right-side regions are even (e.g., 12, 24, 34, …) and all left-side regions are odd (e.g., 11, 21, 31, …).

Both atlases are based on ex vivo acquisitions, while our MRI data were acquired in vivo. This mismatch may introduce geometric distortions due to tissue shrinkage and altered contrast in the ex vivo templates, potentially affecting registration accuracy and regional quantification, especially in peripheral or ventricular regions. To minimize these effects, non-linear registration algorithms were applied.

Each image was segmented into grey matter, white matter and cerebrospinal fluid based on the SIGMA tissue probability maps. When necessary, intensity modulation was applied to preserve original tissue volumes in the registered images. Regional volumes and tissue percentages were calculated by applying the SIGMA atlas to the individual grey and white matter segmentations. To obtain tissue volume and percentages in the Waxholm atlas space, linear registration from the SIGMA space to the Waxholm space was performed using ANTsPy (v. 0.5.3).

For voxel-based morphometry, further normalization of the segmentation to a DARTEL template was performed. The template was generated using animals from each group (4 from each group, pre-and post-intervention). Images were smoothed with a 6 × 6 × 6 mm^3^ kernel (equivalent to 0.6 × 0.6 × 0.6 mm^3^ for the original image size) before comparison with the general linear models.

The SIGMA atlas, originally based on Wistar rats, was used for cross-validation, while the WHS atlas (Sprague-Dawley) was prioritized for primary analyses to match our animal strain.

### Image analysis

Regional grey matter density (GMD, %) and volume (cc) data were extracted for each rat and each brain region, separately for the SIGMA and WHS atlases. GMD was defined as the percentage of grey matter tissue within each region of each hemisphere, as determined by segmentation of T2*-weighted images using the SIGMA tissue probability maps^16^. GMD estimated from T2*-weighted MRI and atlas-based segmentation is an indirect measure reflecting the proportion of voxels classified as grey matter within a given anatomical region, based on intensity-based tissue probability mapping. While this metric is widely used in preclinical imaging to explore regional differences and longitudinal changes in grey matter integrity, its precise cellular correlates, such as neuronal and glial densities or neuropil content, remain incompletely characterized and can be influenced by factors such as partial volume effects, segmentation approach, and tissue properties^32,33^.

### Quantification and Analysis of Measurement Variability

To maximize statistical power and ensure robust reliability assessment, our laboratory leveraged additional imaging data from a distinct cohort initially enrolled in an inflammatory pain model. Although this cohort failed to display the expected inflammatory pain phenotype, all animals underwent the same baseline (T0) brain MRI acquisition. By combining these baseline scans with those from the SNI and healthy control groups, the resulting dataset comprised 30 animals, representing an exceptional sample size for preclinical neuroimaging studies, where typical cohort sizes are limited by feasibility constraints.

To assess the reliability of our segmentation and measurement pipeline, the coefficient of variation (CV) was computed for both GMD and volume across all animals (*n* = 30) at baseline for each hemisphere and region:

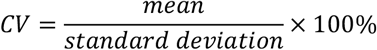

To investigate potential sources of variability, we evaluated the relationship between the CV of GMD and the CV of volume across regions. This analysis aimed to determine whether variability in tissue density measurements was driven by volumetric instability (e.g., registration errors, partial volume effects) or reflected intrinsic biological or technical noise. For this purpose, we generated scatterplots comparing GMD CV (%) against volume CV (%) for all regions within each atlas (see Results, Fig. 3). A robust linear regression was performed to quantify the association, with Pearson’s correlation coefficient used to assess significance (α = 0.05, Benjamini-Hochberg false discovery rate (FDR)-corrected). Regions exhibiting high variability (CV > 50%) were further examined for anatomical or technical peculiarities (e.g., proximity to ventricles, CSF pulsation artifacts, or inconsistent atlas alignment).

### Assessment of Hemispheric Asymmetries

To evaluate potential anatomical asymmetries, we compared GMD values between the left and right hemispheres for each region using paired t-tests. P-values were corrected for multiple comparisons using the FDR procedure (q < 0.05). Based on these comparisons, all subsequent analyses were performed separately for each hemisphere to ensure accurate characterization of regional grey matter density and to preserve sensitivity to potential lateralized phenomena. (see Results section)

### Exclusion Criteria

To ensure the accuracy and reliability of our analyses, several exclusion criteria were systematically applied:

1. **White Matter Regions**: All white matter regions were excluded a priori, as our analyses focused exclusively on grey matter density and volume.
2. **Incomplete Brain Coverage**: Due to technical limitations inherent to our imaging setup, it was not possible to acquire complete brain coverage in all rats. As a result, several regions located at the brain’s anterior (e.g., olfactory bulb and associated layers) and posterior poles (e.g., molecular and granule cell layers of the cerebellum) were only partially included in the acquired images. This incomplete coverage led to artifacts in regional volume estimation and marked dispersion in GMD values for these areas. Accordingly, these regions were systematically excluded from subsequent analyses.
3. **Regions with High Volume Variability and Susceptibility to Partial Volume Effects**: Regions with a coefficient of variation (CV) for volume greater than 50% were excluded from further analysis, as high variability is indicative of unreliable segmentation and partial volume effects, especially in small or anatomically complex regions. For the WHS atlas, most of these excluded regions corresponded to small thalamic nuclei, which also exhibited high variability in GMD, further suggesting partial volume contamination. Using the Siibra 3D atlas tool (EBRAINS)^34^, we confirmed that these regions were primarily adjacent to white matter tracts and/or the ventricles, increasing their susceptibility to partial volume effects. Notably, all excluded regions except the angular thalamic nucleus could be explained by these anatomical considerations; the high variability observed for the angular thalamic nucleus could not be attributed to proximity to white matter or ventricles. For the SIGMA atlas, only three regions were excluded due to a CV for volume exceeding 50%: the pretectal region (adjacent to the ventricles), the associative temporal cortex (at the edge of the field of view), and the dysgranular insular cortex (peripheral location, likely outside the field of view).
4. **Regions with Missing GMD Data**: Any brain regions for which GMD data were missing were also excluded from all analyses to ensure data completeness and integrity.
5. **Composite Regions in the SIGMA Atlas**: To avoid redundancy, composite regions that encompassed multiple subregions were excluded from the SIGMA atlas analyses. For example, the “Primary Visual Cortex” region, which includes both the “Primary Visual

Cortex Binocular Area” and the “Primary Visual Cortex Monocular Area,” was excluded to prevent overlap with these more specific subregions.

These exclusion criteria were implemented to improve the reliability and biological interpretability of our regional grey matter analyses. After all exclusions, 57% (134/234) of SIGMA regions and 58% (256/444) of WHS regions were retained for analysis. The precise anatomical locations and justifications for excluded regions are provided in Supplementary Table 1.

### Longitudinal analysis of Grey Matter Density Changes

To assess longitudinal changes in grey matter density (GMD), we compared pre-intervention (baseline, T0) and post-intervention (28 days after the start of the experiment, T28) measurements for all animals. For each region of interest (ROI), GMD values were extracted at both time points for each subject in both the SNI (n = 9) and control (n = 6) groups.

To provide a comprehensive overview of regional changes across the brain, we calculated the difference in GMD (ΔGMD = GMD_post – GMD_pre) for each ROI and animal and statistical significance was determined for each ROI, and significant differences (p < 0.05, FDR-corrected).

All data processing and statistical analyses were performed using Python (v. 3.8), with specific packages including NumPy, SciPy, pandas, and statsmodels. Visualization and quality control were conducted in 3D Slicer and ITK-SNAP, with additional plots generated using plotly.

### Statistical analysis

For behavioral tests, statistical analysis was performed using GraphPad Prism 7.01 (GraphPad Software, Inc. La Jolla, CA). for imaging, statistical analysis was performed using Python (v. 3.11.5), with specific packages including scipy.stats and statsmodels.stats.multitest. Data were plotted as means ± SEM for all curves and bar graphs.

## Results

### Pain-related behavior evaluation

#### Gray matter densities

To validate the reliability of our imaging and segmentation pipeline, we first quantified the coefficient of variation (CV) of grey matter density (GMD) for each brain region at baseline (n = 30 rats). Across all regions, the mean CV did not exceed 5% (2,27%), indicating minimal variability between subjects and scans at the initial time point. This low level of measurement dispersion demonstrates the high reproducibility and stability of our acquisition protocol, supporting its suitability for detecting biologically meaningful longitudinal changes in subsequent analyses.

Initial analyses revealed that a majority of brain regions exhibited significant hemispheric differences in GMD after correction for multiple comparisons (120 out of 134 regions in the SIGMA atlas and 234 out of 256 regions in the WHS atlas). Given the widespread nature of these asymmetries, pooling left and right hemisphere data would risk obscuring meaningful anatomical or biological effects. Therefore, all subsequent analyses were conducted separately for each hemisphere to ensure accurate characterization of regional grey matter density and to preserve the sensitivity to potential lateralized phenomena. See supplementary Table 2 for GMD interhemispheric results.

#### Comparison of Variability Across Atlases

Scatterplots of GMD coefficient of variation (CV) versus volume CV were generated for all regions in both the SIGMA and WHS atlases (Figure 2). For the SIGMA atlas, the majority of regions displayed low CVs (<10%) for GMD metric, indicating stable and reproducible segmentations across animals. A subset of regions exhibited notably higher variability, clustering predominantly near anatomical boundaries or areas prone to partial volume effects. Pearson’s correlation analysis demonstrated a significant association between volume CV and GMD CV, suggesting that local segmentation instability contributed to measurement dispersion.

**Fig 1.**
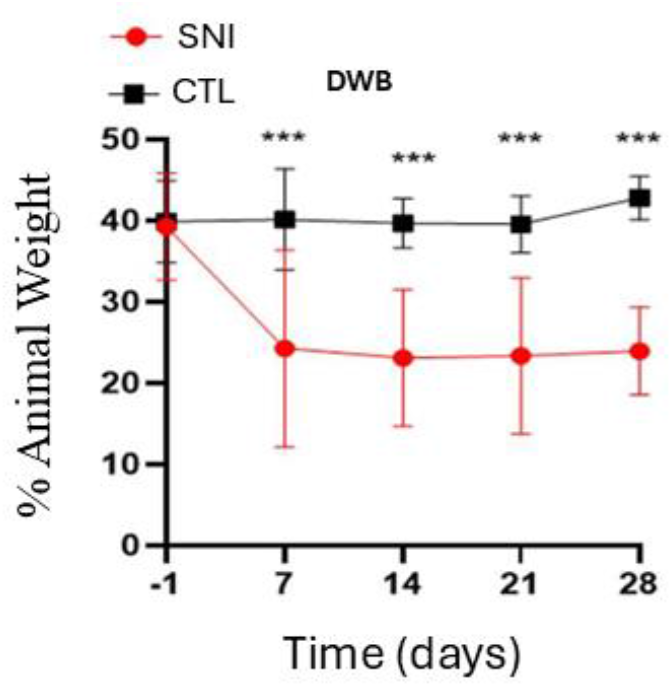
Behavioral outcomes in SNI and control rats for Weight distribution. Rats in the SNI group exhibited a significant decrease in weight distribution on the operated (ipsilateral) hind paw compared to controls by day 7 post-surgery. Data are presented as mean ± SEM. ***p<0.001, **p<0.01, *p<0.05 vs. control (two-way ANOVA with post-hoc test). Gray matter densities

**Figure 2.**
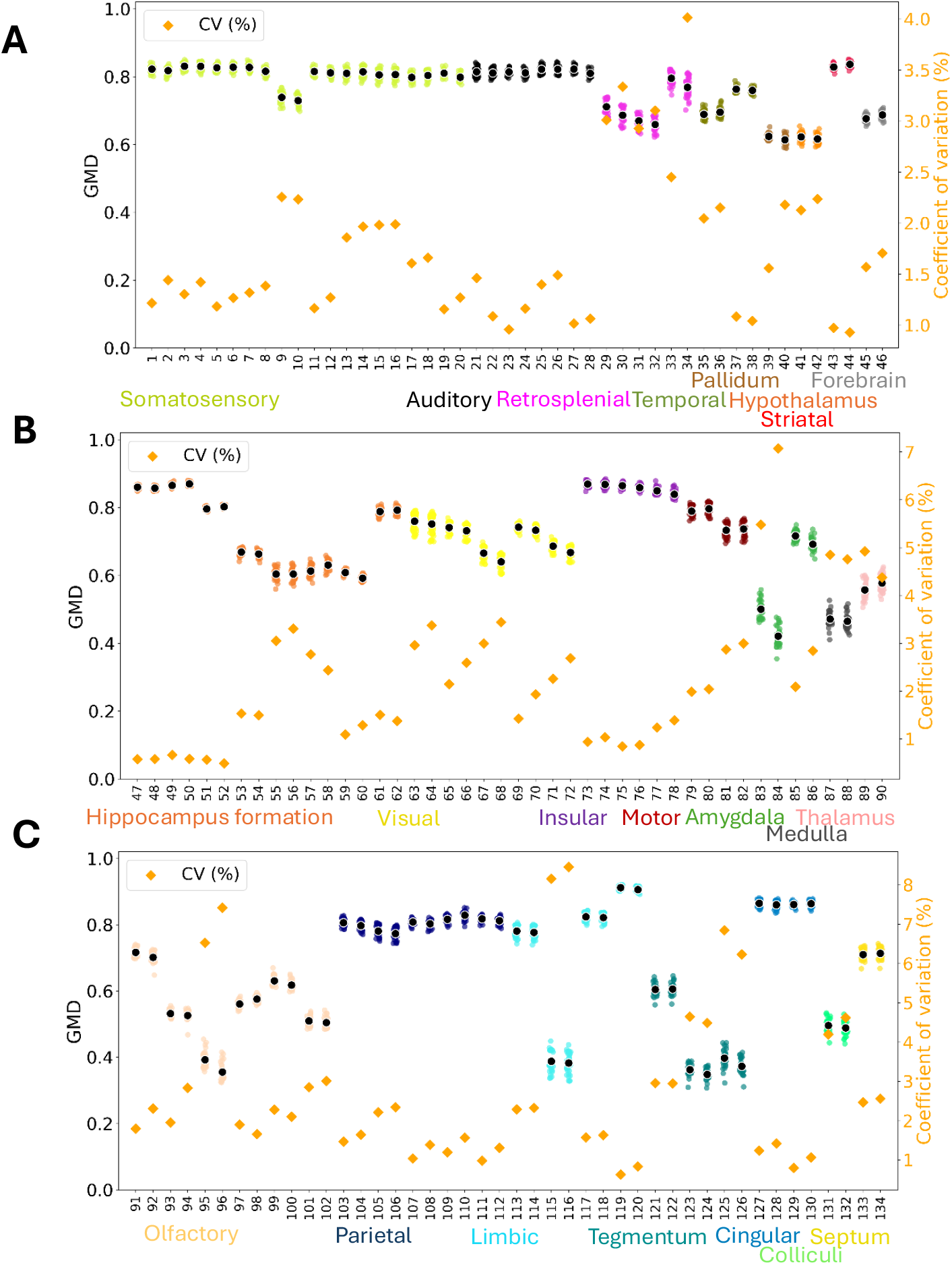
Baseline regional gray matter density (GMD) and coefficient of variation (CV) across all brain regions (n = 30 rats) in SIGMA atlas. On each subpanel, the left y-axis indicates gray matter density (GMD, scale: 0 to 1), while the right y-axis shows coefficient of variation (CV, %). (A) GMD and CV of Somatosensory, Auditory, Retrosplenial, Temporal and Striatal System and Pallidum, Hypothalamus and Forebrain. (B) GMD and CV of Hippocampus formation, Visual, Insular and Motor systems, Amygdala, Medulla and Thalamus. (C) GMD and CV of Olfactory, Parietal, Limbic and Cingular systems, Tegmentum, Colliculi and Septum. Numbers represent the regions, see supplementary Table 1 for correspondence.

**Figure 3.**
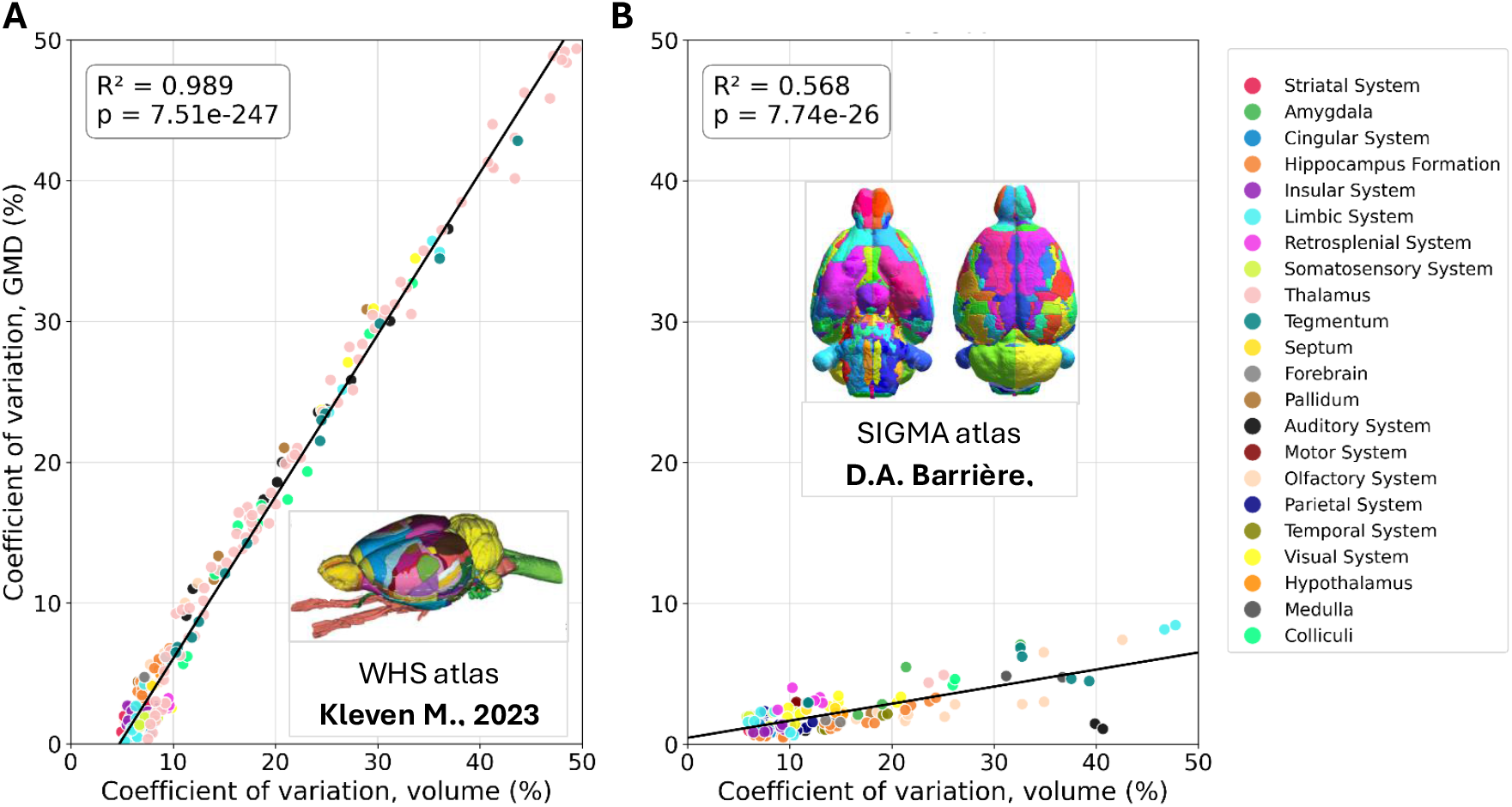
Comparison of Volume and Grey Matter Density Variability Across the SIGMA and WHS Rat Brain Atlases. (A) Waxholm atlas (n = 256 regions); (B) SIGMA atlas (n = 134 regions). Each point = one region; n = 30 rats.

After applying exclusion criteria, regions with volume CV > 50% were systematically excluded from subsequent analyses. The SIGMA atlas retained 134 regions while the WHS atlas retained 256 regions for final analysis. Full lists of retained and excluded regions are presented in Supplementary Table 1.

#### Relationship between Volume and Grey Matter Density Variability Across Atlases

The relationship between volume variability and GMD variability was assessed for all regions and rats using both the SIGMA and WHS atlases (Figure 2A–B). For the SIGMA atlas (Figure 2A), a moderately positive but relatively flat linear relationship was observed between the coefficient of variation (CV) for volume and for GMD (R^2^=0.568, *p*=7.74 × 10^−26^), indicating that in this atlas, regional variability in GMD is only partially explained by volume variability. In contrast, the WHS atlas (Figure 2B) exhibited a quasi-quadratic (x^2^-shaped) association (R^2^=0.989, *p*=7.51 × 10^−247^), whereby GMD variability increased sharply in regions with high volume variability.

Based on these findings, the SIGMA atlas was selected for all subsequent analyses, as the relatively weak dependency between GMD and volume variability suggests greater robustness to segmentation or registration artifacts, likely due to larger region sizes and less stringency in ROI boundaries compared to the WHS atlas.

#### Regional Changes in Grey Matter Density Over Time

Changes in GMD over time were analyzed for both the SNI and control groups (Figure 4). Longitudinal GMD measurements revealed distinct, region-specific effects between SNI and control animals (Figure 4). Scatter plots for six regions (three anatomical hubs, bilateral) illustrate these changes. In the Frontal Association Cortex (regions #115 and 116) and Primary Cingular Cortex (regions #127 and 128), SNI animals showed a significant increase in GMD from baseline to 28 days in both hemispheres (~10% increase for Frontal Association Cortex, ~2% increase for Primary Cingular Cortex; *p* < 0.05, FDR-corrected), while control animals did not exhibit significant changes over time. In the Amygdalohyppocampic Area (regions #83 and 84), SNI animals exhibited a ~10% decrease in GMD (*p* < 0.05, FDR-corrected), whereas controls showed no significant change. These data highlight the specificity of longitudinal GMD alterations in the context of neuropathic pain.

**Fig. 4.**
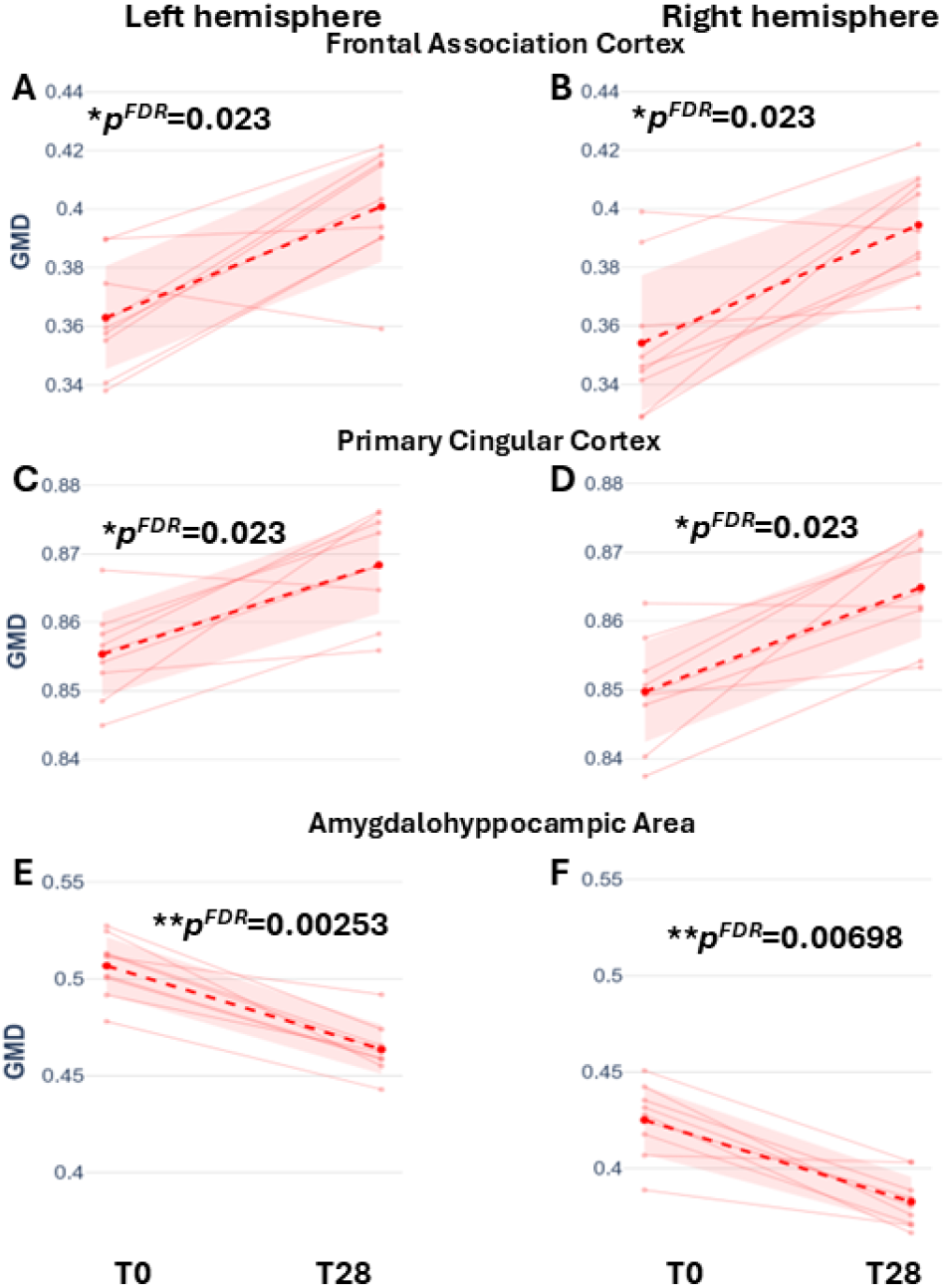
Longitudinal Changes in Grey Matter Density in Corticolimbic Regions in SNI group. Scatter plots depict individual GMD values at baseline (T0) and 28 days post-intervention (T28) in both hemispheres for (A, B) Frontal Association Cortex, (C, D) Primary Cingular cortex, and (E, F) Amygdalohyppocampic, illustrating SNI group changes. *\*\*p<0*.*01, *p<0*.*05*. Results for the control group (n=6) did not reach statistical significance and are provided in Supplementary Table 1 for reference.

All other regions are summarized globally in overview bar plots (Figure 5) and supplementary tables 1.

**Figure 5.**
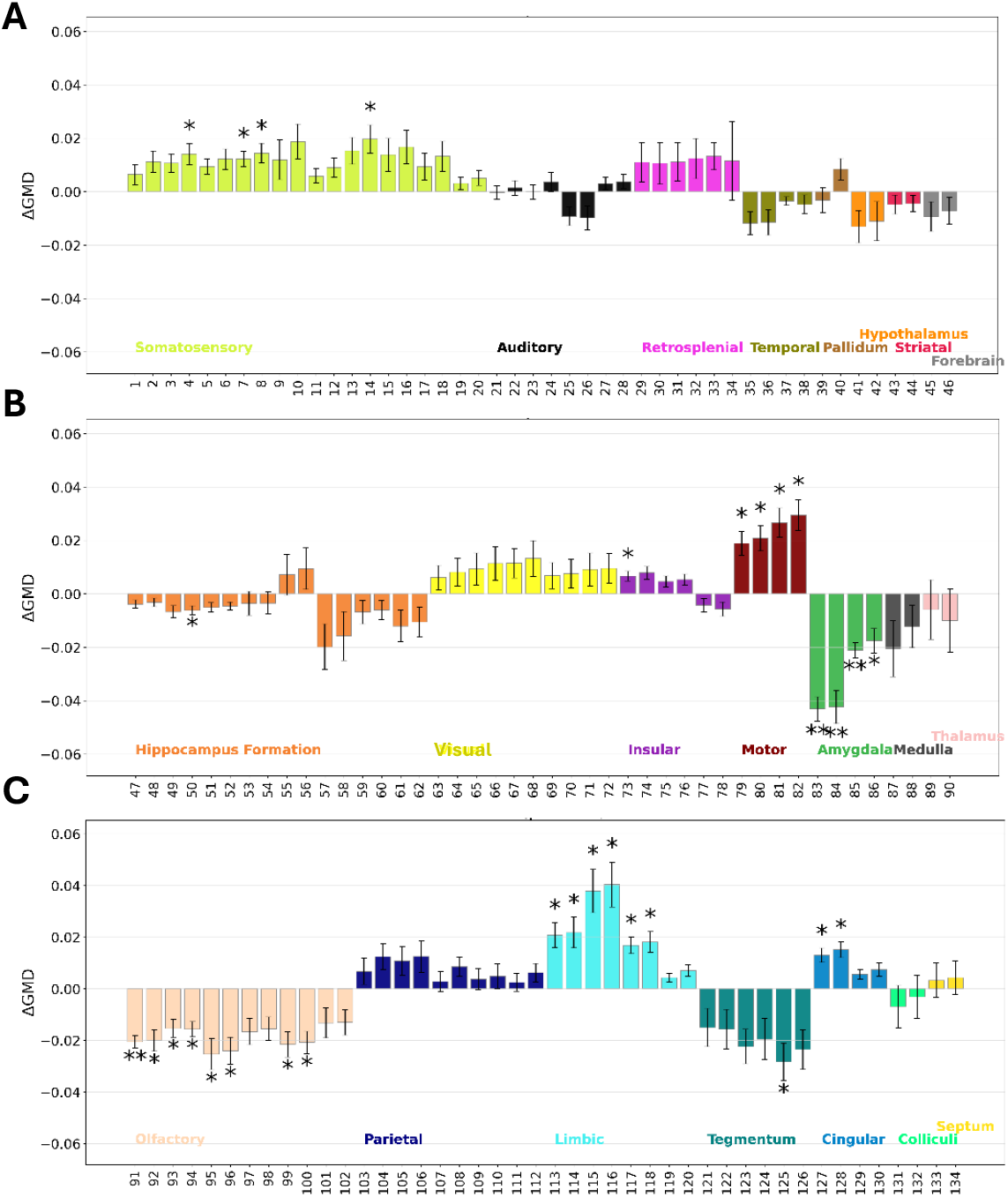
Regional Changes in Grey Matter Density in SNI Rats, Grouped by Anatomical System. Mean ΔGMD per brain region is shown and grouped by major anatomical system. Each bar displays the difference between T0 and T28 (mean ± SD) for the SNI group. Asterisks indicate regions with significant GMD change following FDR correction (*p<0.05, **p<0.01). (A) Mean ΔGMD and CV of Somatosensory, Auditory, Retrosplenial, Temporal and Striatal System and Pallidum, Hypothalamus and Forebrain. (B) Mean ΔGMD and CV of Hippocampus formation, Visual, Insular and Motor systems, Amygdala, Medulla and Thalamus. (C) Mean ΔGMD and CV of Olfactory, Parietal, Limbic and Cingular systems, Tegmentum, Colliculi and Septum. Numbers below/above each bar correspond to the specific region; the full correspondence between region number and anatomical name is provided in Supplementary Table 1.

Out of the 134 retained regions, 31 regions exhibited significant GMD changes in SNI animals, predominantly within the limbic and prefrontal circuitry, whereas auditory, parietal, visual, retrosplenial, and temporal cortices remained unchanged. In total, 21% of the mean brain volume in SNI rats showed significant GMD changes, of which over 17% underwent an increase in GMD and ~3% underwent a decrease. No significant changes in GMD were detected in the control group, which is why only SNI group results are illustrated here. The results for both groups, including all statistics for control animals, can be found in Supplementary Table 1.

## Discussion

In this study, dynamic weight bearing (DWB) analysis confirmed the development of chronic pain in SNI rats^35^, as evidenced by a significant and sustained decrease in weight distribution on the operated hind paw compared to controls from day 7 post-surgery onward. These findings are consistent with established literature demonstrating that nerve injury leads to persistent functional deficits and spontaneous pain behaviors in rodent models of neuropathic pain^28,35,36^. The marked difference between SNI and healthy groups validates the utility of the DWB as an objective metric for chronic pain, providing a robust behavioral phenotype upon which to base subsequent analyses^37^. Importantly, this confirmation enables the present study to concentrate on exploring the neuroanatomical and neuroimaging correlates of chronic pain, confident that the behavioral manifestation of neuropathy has been reliably established in the experimental cohort.

Our longitudinal analysis revealed a distinct pattern of regionally specific gray matter density (GMD) changes in SNI rats, aligning with and extending key concepts outlined in the introduction. As hypothesized from human neuroimaging findings, the most prominent changes occurred within the limbic, prefrontal, and cingulate circuits networks repeatedly implicated in the chronification and emotional modulation of pain^1,9–11,38,39^. Notably, approximately 21% of total brain volume showed significant GMD alterations in the SNI group, with the majority (about 17%) reflecting increases and a smaller fraction (about 3%) showing decreases, emphasizing both the extent and heterogeneity of neuroplastic responses to chronic pain.

Strikingly, GMD increases were most pronounced in regions such as the frontal association cortex, cingulate cortex, and select entorhinal and amygdalar areas. These structures, central to affective and motivational aspects of pain processing, have previously been linked to the transition from acute to chronic pain and the emergence of pain-related comorbidities such as anxiety and depression^1,10,11,38–40^. Our data suggest that the sustained nociceptive input and behavioral phenotype of chronic pain are accompanied by adaptive morphometric remodeling, potentially reflecting enhanced synaptic connectivity, dendritic reorganization, or compensatory mechanisms within these circuits^1,11,39^.

In contrast, no significant changes in GMD were detected in primary sensory, parietal, visual, retrosplenial, or temporal cortices, supporting the notion that as pain becomes chronic, its representation shifts away from classical sensory processing networks toward limbic-motivational regulation^11,15,21,38,39^. This regional divergence directly mirrors the shift in functional brain activity described in longitudinal fMRI and PET studies, where connectivity reallocation supports persistent pain experience and associated affective states^10,13,14,19,20,39^.

These findings reinforce the translational relevance of GMD and volume-based morphometric measures, as highlighted in clinical meta-analyses, for identifying neural signatures and potential therapeutic targets in chronic pain conditions^1,11,38,39^.

By bridging longitudinal MRI data with behavioral and regional analyses, this study clarifies the spatiotemporal dynamics and circuit specificity of pain-induced plasticity^9,10,13,38,39^. These findings pave the way for future investigations into the reversibility, persistence, and functional consequences of morphometric changes in chronic pain, as well as their utility for biomarker and intervention strategies in translational neuroscience^1,11,20,38,39^.

## Limitations

The selection of an appropriate rat brain atlas for morphometric studies remains highly limited, with only a handful of publicly available resources^41^ offering the necessary anatomical annotation and imaging compatibility for robust quantitative analysis. SIGMA and WHS are among the most widely validated and adopted atlases in preclinical neuroimaging, each providing distinct advantages: SIGMA offers macro-anatomical parcellation suitable for global analyses^29^, while WHS delivers fine-grained regional distinction critical for more precised localized investigations^30^. Comparing both atlases in this context was thus a strategic decision to maximize interpretability and relevance, enabling direct evaluation of methodological trade-offs stemming from region granularity and atlas origin. This approach reflects best practice in the field, where standardized options for rat brain mapping are scarce and careful selection is essential to ensure accuracy and comparability of morphometric findings^41^.

The strong correlation between grey matter density (GMD) and regional volume using the WHS atlas suggests that measurements are strongly influenced by technical artifacts such as registration errors, atlas–image mismatches, and partial volume effects, especially in small regions where voxels capture mixed tissue classes. In contrast, with the SIGMA atlas, GMD and volume also covary but with a shallower slope, indicating that GMD is more stable across region sizes and less dominated by partial volume effects, though biological variability also appears modest. Some variability under SIGMA may further reflect the use of a Wistar-derived template rather than Sprague-Dawley anatomy.

These atlas-related trade-offs underscore that coarser parcellations (SIGMA) reduce partial volume noise but may obscure subtle, localized changes, whereas finer atlases (WHS) increase anatomical detail but amplify technical confounds. Validation with complementary methods (e.g., histology, multi-atlas consensus) and further optimization of segmentation and registration remain necessary.

Although no GMD changes were detected in controls, the absence of a sham-operated group limits the ability to fully separate effects of nerve injury from surgical or procedural factors. However, this design choice was deliberate: using intact controls better parallels human neuroimaging studies, where comparison groups typically do not undergo sham interventions. This approach strengthens translational relevance, though it still requires some caution in attributing observed anatomical changes solely to neuropathic pain.

Finally, our findings apply exclusively to male rats; additional studies including females are needed to assess whether similar structural adaptations and atlas performance hold across sexes^42–44^. Therefore, caution is warranted when attributing anatomical changes solely to neuropathic pain or when generalizing beyond the present model and cohort.

## Conclusion

This study demonstrates that high-resolution MRI can robustly and objectively quantify the widespread and regionally specific reorganization of the male rat brain induced by chronic neuropathic pain, echoing findings from other neuroimaging modalities^1,9–11,38,39^Our results confirm that these structural changes are most pronounced within limbic and prefrontal systems, central to pain chronification and emotional comorbidities—while sparing primary sensory and associative cortices^1,10,11,39^. However, the precise cellular and molecular substrates underlying these volumetric and density alterations remain elusive.

Future research must now focus on integrating advanced imaging with cellular and molecular analyses, such as histology, transcriptomics, and circuit-level functional approaches, to disentangle the mechanisms driving pain-induced neuroplasticity^38,39^. In addition, expanding such investigations to include female animals will be essential to better capture sex differences in pain-related brain plasticity and translational relevance^42,44^. Such integration will be crucial to identify targets for prevention or reversal of pathological brain reorganization, and to guide therapeutic strategies that could modify or restore neural network stability in chronic pain^10,20,38,39^. Ultimately, this line of research holds promise for the development of translational biomarkers and personalized treatment options by bridging large-scale MRI phenotypes with molecular and cellular profiles^1,10,11,38,39^.

## Code availability

The codes generated during the current study for the processing of the images are fully available from the GitHub platform using the following link: https://github.com/Tetreault-Pain-Imaging-Lab

## Competing interests

The authors declare no competing interests.

### Supplementary information

Supplementary table 1.csv

Supplementary table 2.csv

## Notes

https://github.com/Tetreault-Pain-Imaging-Lab

### Competing Interest Statement

The authors have declared no competing interest.

### Funding

- Natural Sciences and Engineering Research Council: RGPIN-2021-03669

